# Menstrual cycle phase alters corticospinal excitability and spike-timing-dependent plasticity in healthy females

**DOI:** 10.1101/2025.09.30.679456

**Authors:** P. Spillane, E. Pastorio, E. Nédélec, J. Piasecki, S. Goodall, K M. Hicks, P. Ansdell

## Abstract

The known fluctuations in ovarian hormone concentrations across the eumenorrheic menstrual cycle contribute to modulations in cortical excitability and inhibition. However, how such changes affect spike-timing-dependent plasticity (STDP) has not been systematically studied. This research aimed to determine the effect of the menstrual cycle on corticospinal excitability and STDP.

Twelve eumenorrheic female participants (age: 25 ± 5 years), visited the lab in three menstrual cycle phases: early follicular (EF), late follicular (LF), and mid-luteal (ML). Visits comprised of corticospinal excitability (motor evoked potential [MEP]/M_max_), short-intracortical inhibition (SICI), and intracortical facilitation (ICF) measures, recorded in the resting *first dorsal interosseous*. Followed by a paired associative stimulation (PAS) protocol, utilising ulnar nerve and transcranial magnetic stimulation (25 ms interstimulus interval) to elicit neuroplasticity. To assess the time course of STDP, measurements were repeated at 15 and 30-minutes post PAS.

Corticospinal excitability (MEP/M_max_) was greater in the LF phase (*p*≤0.002) compared to EF and ML, with no phase effects observed for SICI or ICF (*p*≥0.112). PAS elicited an increase in MEP/M_max_ across all phases at 15-minutes (112 ± 5, 115 ± 5, and 113 ± 7% baseline, *p*≤0.010), whereas at 30-minutes only ML was facilitated (126 ± 7% baseline, *p*=0.029).

The present data demonstrates facilitatory STDP can be induced with PAS across the tested menstrual cycle phases, but responses are prolonged and potentiated in the ML phase. Additionally, increased corticospinal excitability in the LF phase is likely due to intrinsic changes within the descending tract, as no changes in intracortical neurotransmission were observed.

## Introduction

There is a significant sex gap in scientific research whereby females have often been excluded in biological studies, including neurophysiology (Jenz & Pearcey, 2022). Circulating ovarian hormone levels routinely fluctuate across the menstrual cycle (Malcolm & Cumming, 2003). Concerns about this additional variability are often cited as a barrier to including females in such research (Woitowich et al., 2020), despite these cyclical changes being experienced by ∼50% of females in the United Kingdom (ONS, 2023). Although female inclusion in neuroscience has improved over recent years, this has not been reflected by increased sex comparisons (Woitowich et al., 2020), nor rigorous investigation of the menstrual cycle.

The role of oestrogens and progesterone on excitability of the primary motor cortex and the corticospinal-motoneuronal pathway has previously been studied non-invasively in humans using transcranial magnetic stimulation (TMS; Ansdell et al., 2019; Badawy et al., 2013; Hattemer et al., 2007; Inghilleri et al., 2004; Smith et al., 2002, 2003). The limited available data appears to show that corticospinal excitability remains constant when probed with single pulse TMS (Ansdell et al., 2019). Whereas, conclusions about motor cortical neurotransmission are equivocal, with both intracortical facilitation (ICF) and short-interval intracortical inhibition (SICI) have been identified to fluctuate across the cycle (Ansdell et al., 2019; Smith et al., 2002, 2003), or remain unaffected (El-Sayes et al., 2019; Zoghi et al., 2015). SICI is sensitive to changes in GABA_A_ neurotransmission (Di Lazzaro et al., 2007), which is potentiated by progesterone (Smith et al., 1987a). However, the exact mechanisms behind ICF are less clear, but are likely mediated by glutamatergic neurotransmission (Rossini et al., 2015). Oestrogen promotes the release of, and sensitivity to, glutamate via upregulating the sensitivity and expression of *N*-methyl-D-aspartic acid (NMDA) receptors, which leads to increased neuronal excitability (Adams et al., 2004; Smith et al., 1987b). Collectively, fluctuations in ovarian hormones provide evidence for the acute changes in neurotransmission across the menstrual cycle, yet there is considerably less evidence about how these changes might modulate neuroplasticity.

TMS can also be used to modulate excitability within the motor cortex, either in a non-selective manner using repetitive TMS (rTMS; Jannati et al., 2022) or selectively on synapses utilising forms of spike-timing-dependent plasticity (STDP) such as paired associative stimulation (PAS; Stefan et al., 2002). In females, facilitatory responses to PAS neuroplasticity protocols are blunted in older adults, which has been linked to the decline in sex hormone levels after the menopause (Polimanti et al., 2016; Tecchio et al., 2008). This reduction is thought to be a response to a loss of oestradiol blunting long term potentiation-like plasticity (LTP), mediated through lower NMDA receptor transmission (Smith & McMahon, 2005). The effects of the menstrual cycle on cortical neuroplasticity have been further investigated using two different rTMS paradigms, low frequency rTMS (Inghilleri et al., 2004) and intermittent theta burst stimulation (iTBS; Ramdeo et al., 2024). Although each assessed only two menstrual cycle phases, both reported that elevated oestrogen during the follicular phase facilitates LTP-like neuroplasticity, whilst responses in the luteal phase were blunted. There were also discrepancies in the tested phases and the methods of characterising the menstrual cycle, with Ramdeo et al. (2024) using ovulation prediction, and Inghilleri et al. (2004) relying on calendar counting. As yet, no one has applied the three-step menstrual cycle phase verification method (Schaumberg et al., 2017) to comprehensively investigate the impact of the menstrual cycle on STDP.

As well as being a selective manner of inducing LTP, PAS has been shown to elicit a facilitatory response in a greater proportion of people than the non-focalised rTMS protocols (Player et al., 2012), making it a more useful tool for probing neuroplastic capacity. Some of this variation might be explained by stimulation intensity, as typically rTMS (including iTBS) protocols use subthreshold stimuli (Jannati et al., 2022; Maeda et al., 2000), whilst PAS utilises a suprathreshold stimulus (Suppa et al., 2017), which gives confirmation of stimulation site and intensity throughout the protocol. Therefore, the selective and consistent nature of neuroplasticity induced by PAS and other STDP protocols has important applications in neurorehabilitation (Grover et al., 2023; Suppa et al., 2017). For example, priming with PAS has the potential to modulate motor skill learning outcomes (Jung & Ziemann, 2009). Combined with best practice methods of menstrual cycle research, PAS offers a novel method to investigate the influence of sex hormones on cortical synaptic neuroplasticity, which could begin to optimise neurorehabilitation protocols according to sex hormone concentrations.

Accordingly, the aim of this study was to assess the effect of menstrual cycle phase on motor cortical inhibition and facilitation, corticospinal tract excitability, and STDP. It was hypothesised that the high oestrogen state during the late follicular phase would result in greater neuroplasticity and motor pathway excitability compared to the early follicular phase. It was also hypothesised that these changes would be nullified in the inhibitory presence of progesterone during the mid-luteal phase.

## Methods

### Ethical Approval

The present study received institutional ethical approval from the Northumbria University Health and Life Sciences Research Ethics Committee (reference: 2878) and was conducted according to the Declaration of Helsinki in all aspects apart from registration in a database. All participants gave their written informed consent prior to any part of the study.

### Sample Size Estimation

Sample size was estimated from the F statistic from Inghilleri et al. (2004) for the three-way interaction between menstrual cycle phase, stimulation number, and sex in rTMS pulse trains (F_9,126_ = 3.44). This was then converted to an effect size as per Lakens (2013*;η*p*²* = 0.197). With the parameters of *α* = 0.05 and 1-*β* = 0.95, the minimum sample size required was 13 participants. Therefore, to maximise statistical power and to account for potential drop out and anovulatory cycles, 28 participants were initially recruited.

### Participants

A total of 28 healthy cis-gender females (age: 25 ± 5 years), who self-reported a regular menstrual cycle (≥21 and ≤35 days) volunteered for the study. Of these, 12 participants completed all three experimental testing sessions and met post-hoc hormonal inclusion criteria (stature: 166.7 ± 5.2 cm, mass: 68.9 ± 8.1 kg, age: 25 ± 5 years). Participants reported as not having taken any form of hormonal contraceptives for at least 6 months prior to participation, were recreationally physically active, self-reporting 435 ± 271 min/week of moderate to vigorous physical activity. Prior to any data collection participants were screened for menstrual cycle irregularities, and for electrical and magnetic stimulation safety (Rossi et al., 2011). Participants were not taking any medication known to affect neurological function, and were free of known neurological illness, and musculoskeletal injury to the relevant limbs. Participants were requested to refrain from strenuous physical activity and alcohol (24 h) and caffeine (12 h) prior to each experimental testing session.

### Experimental Design

This study was repeated measures in design with participants visiting the laboratory on four occasions. Firstly, a familiarisation visit, followed by three experimental visits each lasting ∼2 h. Each trial occurred at the same time of day (± 1.5 h) in three different phases (early follicular [EF], late follicular [LF], and mid-luteal [ML]) of the menstrual cycle, that represented the most distinct hormonal concentrations (Elliott-Sale et al., 2021). Prior to data collection participants were asked to calendar track their menstrual cycle for at least one month, and use urinary luteinising hormone (LH) surge detection kits from the day after menses ceased until the day of LH surge to predict ovulatory status. This first tracked cycle was used to individualise the schedule for the subsequent testing days. Urinary LH detection kits were also used to indicate probable ovulation during the experimental cycle(s), and blood samples collected to retrospectively confirm this via serum hormone concentrations of progesterone. The three phases were defined as EF (day 2-4; see Figure 1 for testing visit scheduling and idealised hormonal state), LF (24-48 h prior to LH surge) and ML (6-8 days post LH surge). The order of testing for menstrual cycle phase was pseudorandomised and counterbalanced between participants.

**Figure 1:**
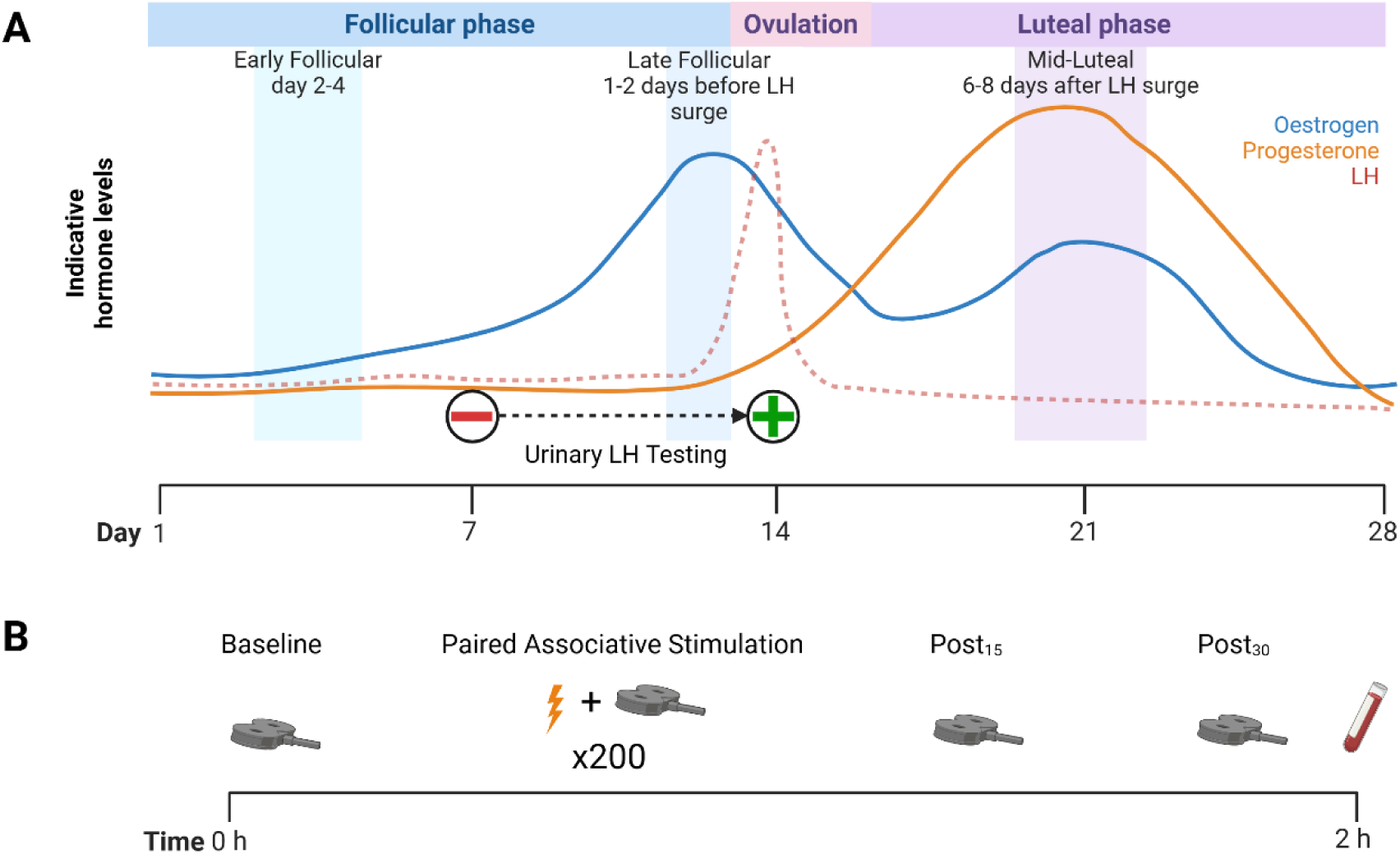
A schematic showing (A) the timing of each lab visit within an idealised menstrual cycle and (B) the structure of each experimental testing session. LH: Luteinising hormone. Created with BioRender.com

The familiarisation visit comprised of introducing participants to the nerve and brain stimulation protocols, including a full baseline assessment and a short (∼2 mins) PAS protocol. The experimental session started with determining stimulation thresholds and intensities at rest in the non-dominant *First Dorsal Interosseous* (FDI) muscle; the maximal compound action potential (M_max_), perceptual threshold, and finally TMS motor threshold (rMT) were recorded. The single and paired pulse TMS assessment was completed at baseline, along with electrical nerve stimulation. Next, the PAS protocol was performed, followed by single and paired pulse TMS assessments 15- and 30-minutes afterwards. Finally, a blood sample was drawn for subsequent hormone analysis. Detailed descriptions of each assessment are provided in the sections below.

### Bipolar Surface Electromyography (EMG)

Surface electromyography was recorded from the FDI on the non-dominant hand throughout assessments of nervous system function and PAS protocol. Hand dominance was determined using the Edinburgh Handedness Inventory (Oldfield, 1971). Two self-adhesive Ag/AgCl recording electrodes (20 × 15 mm, Neuroline 700, Ambu, Denmark) were placed ∼1 cm apart on the belly of the FDI, the reference electrode was placed over the ulna styloid process of the same arm. The raw EMG signal was amplified (×1000), band pass filtered (20 – 2000 Hz) and digitised (5000Hz).

### Electrical Stimulation of the Ulnar Nerve

Activation of the FDI was achieved by percutaneous stimulation of the ulnar nerve; using surface electrodes (3.2 cm diameter; ValuTrode, Axelgaard Manufacturing, Denmark) with the cathode placed proximally to reduce the necessary current (Pieber et al., 2015). The stimulating electrodes were connected to a constant current muscle stimulator (DS7AH, Digitimer, Welwyn Garden City, UK), delivering single pulses of 200 *µ*s duration. During the delivery of electrical stimuli, participants were instructed to keep the target muscle relaxed, confirmed via visual inspection of the EMG trace. For M_max_ assessment stimulation intensity began at 30 mA and was increased by 10 mA until a plateau in peak-to-peak amplitude was observed. To ensure supramaximal stimulation and account for potential changes in axonal excitability (Thomas et al., 2016), the intensity used for all M_max_ assessments was increased by an additional 30%. Both stimulation intensity (EF: 139 ± 51, LF: 148 ± 47, ML: 121 ± 30 mA, F_2,22_ = 2.946, *p* = 0.074) and M_max_ amplitude (EF: 11.92 ± 3.24, LF: 11.65 ± 2.78, ML: 12.06 ± 5.49 mV, F_1.245,13.697_ = 0.540, *p* = 0.869) were consistent across phases. For sensory stimulation, the same electrode placement and stimulus duration was utilised. Determination of perceptual threshold was achieved by starting at an intensity of 1 mA, which was increased in 1 mA increments until participants reported being able to feel the stimulus. This was confirmed by reducing the intensity by one increment, and retesting whether the participants reported being able to feel the stimulus. To ensure effective sensory stimulation, the intensity was then increased to 300% of perceptual threshold (Stefan et al., 2000). Perceptual threshold was constant across the phases tested (4 ± 1 mA for all phases, F_2,22_ = 2.406, *p* = 0.113).

### Transcranial Magnetic Stimulation

Magnetic stimuli were delivered over the motor cortex contralateral to the non-dominant limb, using a flat 70 mm figure of eight coil (Second Generation Remote 3190-00, Peak Magnetic Field 2.2 T) connected to two linked monopulse magnetic stimulators (MagStim BiStim^2^ and 200^2^, The Magstim Company, Whitland, UK). The TMS coil was orientated to induce a posterior-anterior directed magnetic field, delivering a 1 ms pulse. The “hotspot”, defined as the largest and most consistent MEP measured in the FDI, was located at the beginning of each visit, and the site was marked on the scalp with indelible marker pen to ensure consistent coil placement for each subsequent stimulation. rMT was determined according to the IFCN definition (Rossini et al., 2015) as the lowest stimulation intensity (% Maximal Stimulator Output [MSO]) that induced a motor evoked potential (MEP) in the relaxed FDI of >50 *µ*V in 5 out 10 trials and is then increased by 1% MSO. rMT was consistent across the menstrual cycle phases tested (EF: 49 ± 11, LF: 51 ± 12, ML: 50 ± 10% MSO, F_2,22_ = 9.250, *p* = 0.260). This intensity was subsequently used to set the stimulation intensity for measuring corticospinal excitability, which was performed at 120% rMT. The peak-to-peak amplitude of MEPs (mV) was measured in Spike2 software (v10.08, CED, Cambridge, UK) and expressed as a % M_max_. Twenty unconditioned MEPs were recorded to collect a reliable measure of corticospinal excitability (Brownstein et al., 2018). Visual inspection of each trial was performed and any with voluntary activity in the proceeding 100 ms were discarded.

Cortical inhibition was measured using SICI; with a conditioning pulse set at 80% rMT, followed by a test pulse at 120% rMT, with an interstimulus interval (ISI) of 2 ms (Kujirai et al., 1993). Facilitatory neurotransmission was measured using ICF, for which the optimal combination of conditioning stimulus intensity and ISI are undetermined (Brownstein et al., 2018). So for consistency with SICI, and in accordance with studies in the FDI (Zoghi et al., 2015), ICF was quantified using a conditioning stimulation at 80% rMT and a test stimulus at 120% rMT with an ISI of 12 ms (Kujirai et al., 1993). To ensure sufficient data for both SICI and ICF, the amplitude of 20 evoked responses was used in each assessment (Brownstein et al., 2018). Visual inspection of each trial was performed and any with voluntary activity in the proceeding 100 ms were discarded. Paired pulse stimuli were expressed as a percentage of the average unconditioned MEP amplitude at each respective time point.

### Paired Associative Stimulation

The assessment of cortical STDP comprised of a PAS protocol followed by single and paired pulse TMS assessments. LTP-like neuroplasticity was induced using PAS, where the repetitive pairing of a sensory stimulus of a mixed peripheral nerve, with stimulation of the motor cortex causes Hebbian plasticity. The antidromic signal was elicited with an electrical stimulation of the ulnar nerve (see *Electrical Stimulation of the Ulnar Nerve* for set up), the intensity was set at 300% of sensory threshold, aligning with the majority of PAS studies (Carson & Kennedy, 2013). The ISI between sensory stimulus and TMS was set at 25 ms, chosen to elicit a facilitatory PAS response (Stefan et al., 2000). The PAS protocol consisted of 200 paired stimuli delivered at 0.25 Hz. In instances where participants had an rMT above 60% MSO, the frequency was reduced to 0.2 Hz to permit the TMS to recharge between stimuli. The inclusion of frequency within the statistical model did not improve model fit (*p* = 0.956), therefore had no influence on outcomes. Participants were instructed to remain relaxed throughout the protocol, which was confirmed by visual inspection of the electromyogram. The protocol lasted 13 min 20 s at 0.25 Hz, and 16 min 40 s at 0.2 Hz.

The magnitude and time course of neuroplasticity were assessed by repeating the MEP, SICI, and ICF measurements 15 and 30 minutes after PAS, using the protocols described in *Transcranial Magnetic Stimulation*. MEPs were expressed relative to M_max_ at the respective time point (MEP/M_max_), and as a percentage of baseline amplitude. The follow-up assessments tracked the time course of the potentiation effect, with 30 minutes post PAS thought to be the most reliable time point for assessing facilitatory changes in excitability (Alder et al., 2019; Wischnewski & Schutter, 2016).

### Hormone Analysis & Menstrual Cycle Phase Verification

Venous blood samples were collected at the end of each testing session. Each blood draw consisted of a 10 ml sample collected from an antecubital vein, which was left upright for 2 h to coagulate. Samples were centrifuged for 15 minutes at 1000 *g*, and the serum supernatant was then pipetted into aliquots and stored at −80°C until hormone analysis. Hormone analyses were performed at the Bioanalytical Facility, University of East Anglia (Norwich, UK) and undertaken in Good Clinical and Laboratory Practice conditions. The serum were analysed in duplicate for concentrations of 17β-oestradiol and progesterone using electro-chemiluminescence immunoassay on the COBAS e601 automated platform (Roche Diagnostics, Mannheim, Germany). The inter-assay coefficients of variation (CV) were ≤3% within their respective analytical working ranges. The minimum detectable concentrations were 18.4 pmol/L and 0.2 nmol/L for 17β-oestradiol and progesterone, respectively. Hormone levels were used to confirm eumenorrheic status, where a rise in progesterone in the ML phase to a minimum threshold of 9.5 nmol/L was used as the inclusion criterion, indicating probable ovulation (Chinta et al., 2020; Wathen et al., 1984).

### Statistical Analysis

All data processing and analyses were conducted using R Statistical Software (v4.4.1, R Core Team, 2024). Data with a single value in each phase or time point (e.g., hormone concentrations) are presented as mean ± standard deviation (SD) in text, figures and tables. The normality of distribution was confirmed using Shapiro-Wilks tests. For single value baseline variables a one-way repeated measures analysis of variance (1 [variable] × 3 [menstrual cycle phases]) was used to examine phase effects. When sphericity was violated (*p* <0.05) the Greenhouse-Geisser correction was applied to degrees of freedom and significance. Where significance were found, univariate analyses were followed by subsequent pairwise comparisons, with post hoc Tukey correction to identify differences.

For measures that have multiple observations/measurements per time point (e.g., MEP/M_max_) data are presented as estimated marginal means with standard error from model outputs. Linear mixed effects models were used to determine whether variables of interest could be predicted by the fixed effects of menstrual cycle phase for baseline measures; and menstrual cycle phase, time, and their interaction for cortical neuroplasticity, with baseline values included as a covariate (*lme4*, Bates et al., 2015). Random effects included intercepts by participant. Baseline variables model structure was specified as “response ∼ phase + (1 | participant)”. The PAS variables model structure for the added model of main effects was “response ∼ phase + time + baseline response + (1 | participant)”, and interactions as “response ∼ phase * time + baseline response + (1 | participant)”. For PAS measures quantified as percentage of baseline, participant random intercepts were nested with a random slope for menstrual cycle phase (MEP% Baseline ∼ phase * time + (1 phase + | participant). To determine significance likelihood ratio tests (*lmerTest*, Kuznetsova et al., 2017) were performed comparing the full added model against a model without the effect of interest, and interaction model against the full added model. Post hoc pairwise comparisons p-values were adjusted for multiple comparisons using Tukey’s method.

Participant means and phase estimated marginal means were computed for each variable of interest (*emmeans*, Lenth, 2025). Residual plots were visually inspected to confirm homoskedasticity, and where violated data were log transformed for model analysis only. However, data are presented visually and in text following back transformation with an exponential function. Alpha level was set at 0.05 and all data were visualised in R (*ggplot2*, Wickham, 2016).

## Results

### Menstrual Cycle Characteristics

Following screening of 28 potential participants, twelve females were included in the final analyses (stature: 167.6 ± 5.0 cm, mass: 69.8 ± 8.4 kg, age: 25 ± 6 years). A breakdown of the participant attrition is provided in Figure 2, and hormone levels are presented in Figure 3. During the post-hoc hormonal verification, one participant was excluded due to the absence of a rise in 17β-oestradiol at the LF time point and no serum sample at the ML time point, with another failing to exhibit a progesterone rise above 9.5 nmol/L in the ML phase. Both were therefore excluded due to probable anovulatory cycles based on the criteria set out in *Menstrual Cycle Phase Verification* methods section. Five participants started testing in the ML phase, four in the EF, and three in the LF phase. Testing was conducted within one cycle for three participants, two cycles for seven participants, and three cycles for two participants. Visits were conducted on cycle day 3 ± 1 for EF, 13 ± 2 (2 ± 2 days prior to detected LH surge) for those tested in LF phase, and cycle day 20 ± 2 (7 ± 2 days after detected LH surge) for ML. Three participants were tested 1 day post LH surge (1 ± 0 day after detected LH surge) and therefore in the ovulatory phase but data were pooled with the LF phase as hormonal profiles were consistent with the wider sample.

**Figure 2:**
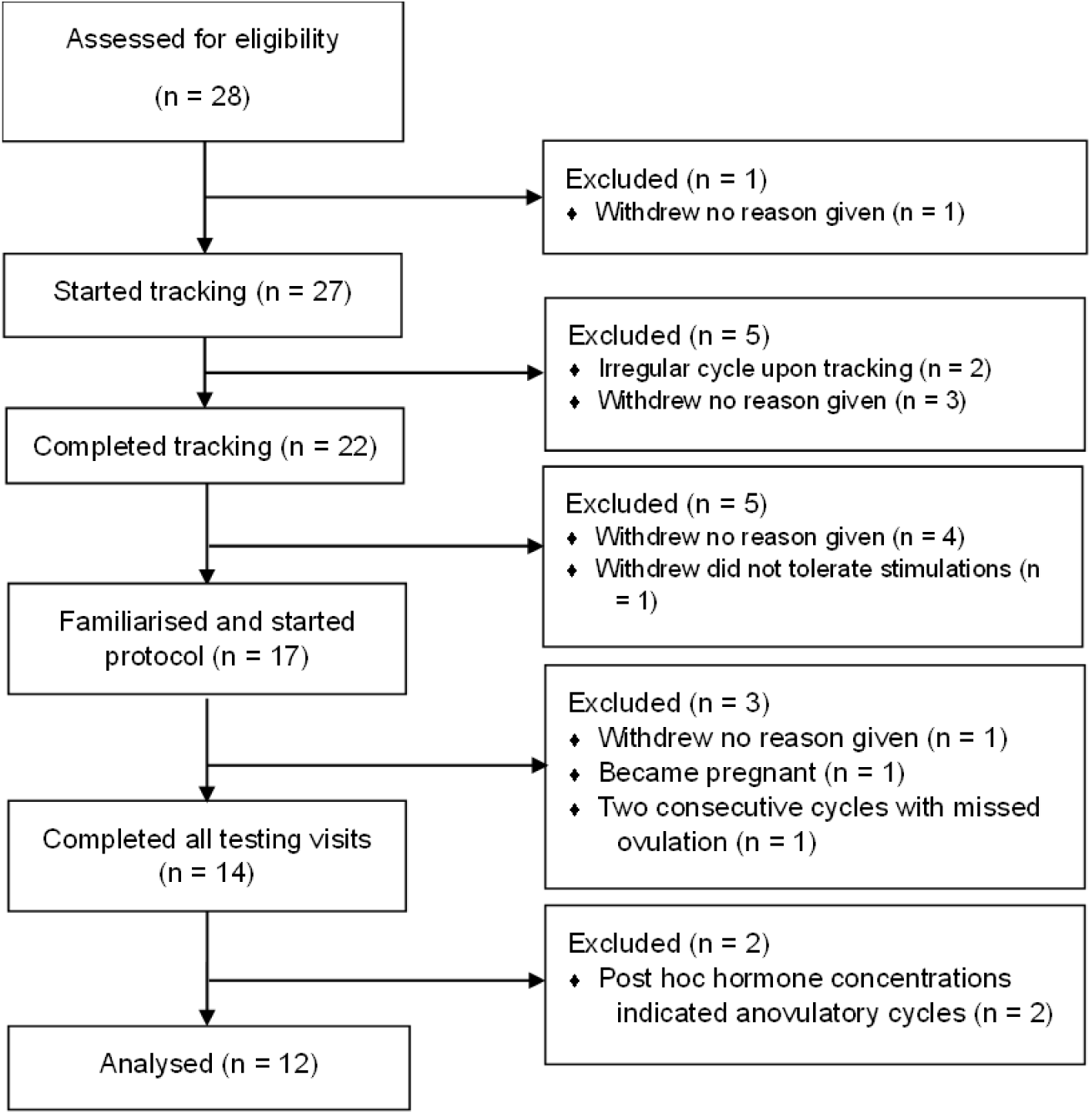
Consort diagram showing the flow of participants from recruitment to completion with the participant attrition and reasons for withdrawal on the right of the diagram.

**Figure 3:**
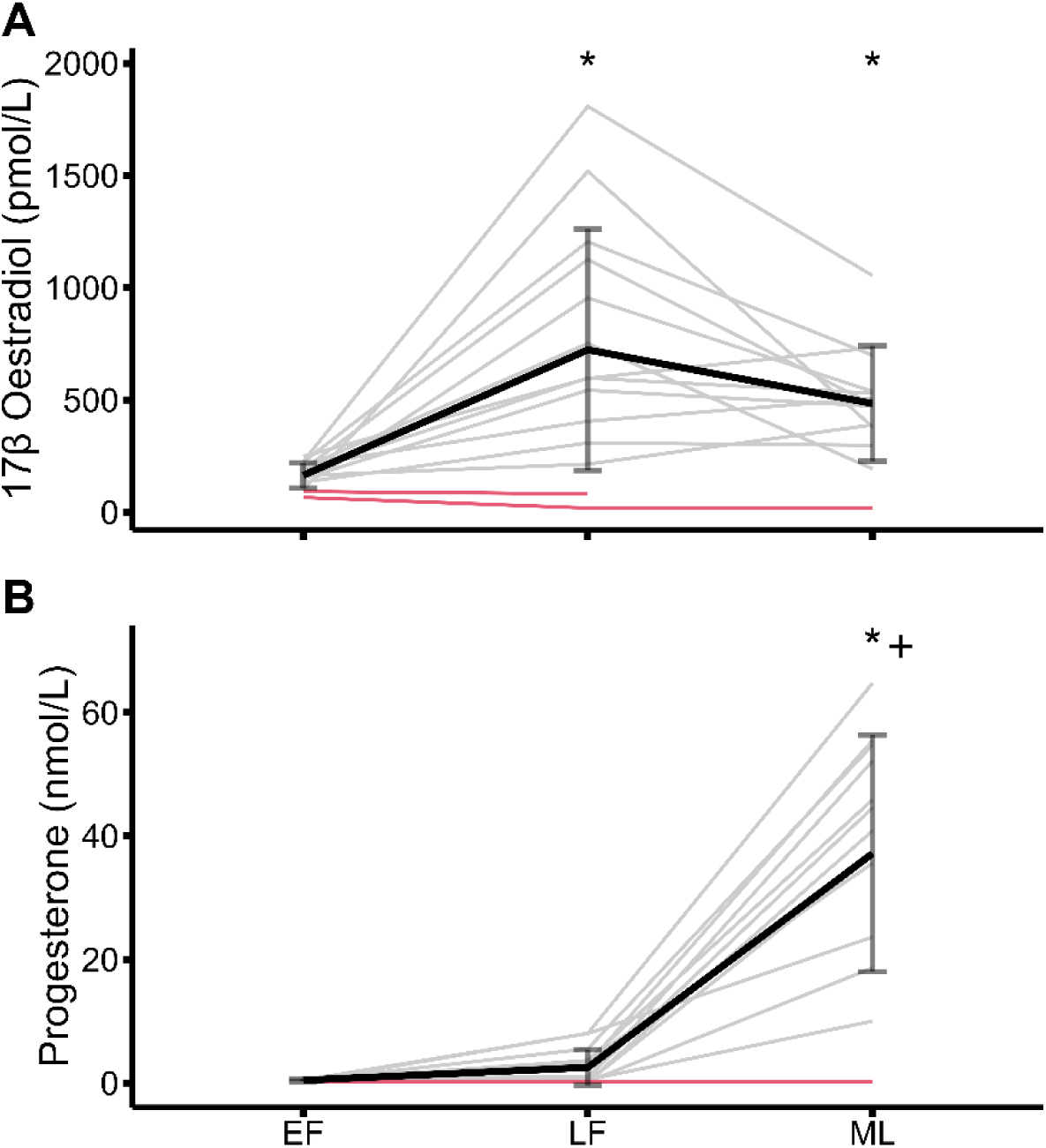
Hormone data for all participants who completed all testing visits. A: Oestradiol concentrations across all three phases. B: Progesterone concentrations across all three phases. Black lines represent group means (error bars indicate SD), grey lines represent participants included for full analysis, whilst red indicates those excluded based on 9.5 nmol/L threshold for progesterone. EF: Early Follicular, LF: Late Follicular, ML: Mid-luteal, * different from early follicular p <0.05. + different from late follicular p <0.05.

As expected, participants displayed large variation in hormone concentrations and patterns across the tested time points (Figure 3). 17β-oestradiol showed a phase effect (F_1.25,15.04_ = 14.76, *p* <0.001), and post hoc tests revealed it to be higher in the LF (835.7 ± 495.8 pmol/L, *p* < 0.001) and ML (522.7 ± 225.5 pmol/L, *p* = 0.003) phases compared to EF (178.2 ± 46.6 pmol/L). Progesterone also demonstrated a phase effect (F_1.03,12.42_ = 46.94, *p* <0.001), with post hoc comparisons being higher in ML (40.3 ± 4.7 nmol/L) phase compared to both EF (0.5 ± 0.2 nmol/L, *p* <0.001) and LF 2.9 ± 3.0 nmol/L, *p* <0.001).

### Baseline neurophysiological assessments

MEP data were logarithmically transformed due to heteroskedastic distribution of the model residuals; however, back-transformed data are presented for clarity. Baseline corticospinal excitability differed across the menstrual cycle (χ^2^ (2)=24.23, *p* <0.001), with the LF (7.7 ± 1.5% M_max_) phase being higher than both EF, (6.2 ± 1.2% M_max_, *p* =0.002) and ML (5.6 ± 1.1% M_max_, *p* <0.001). There was no effect of menstrual cycle phase on SICI (χ^2^ (2)=0.93, *p* 0.627), nor ICF (χ^2^ (2)=4.37, *p* = 0.112). Baseline TMS data are presented in Figure 4.

**Figure 4:**
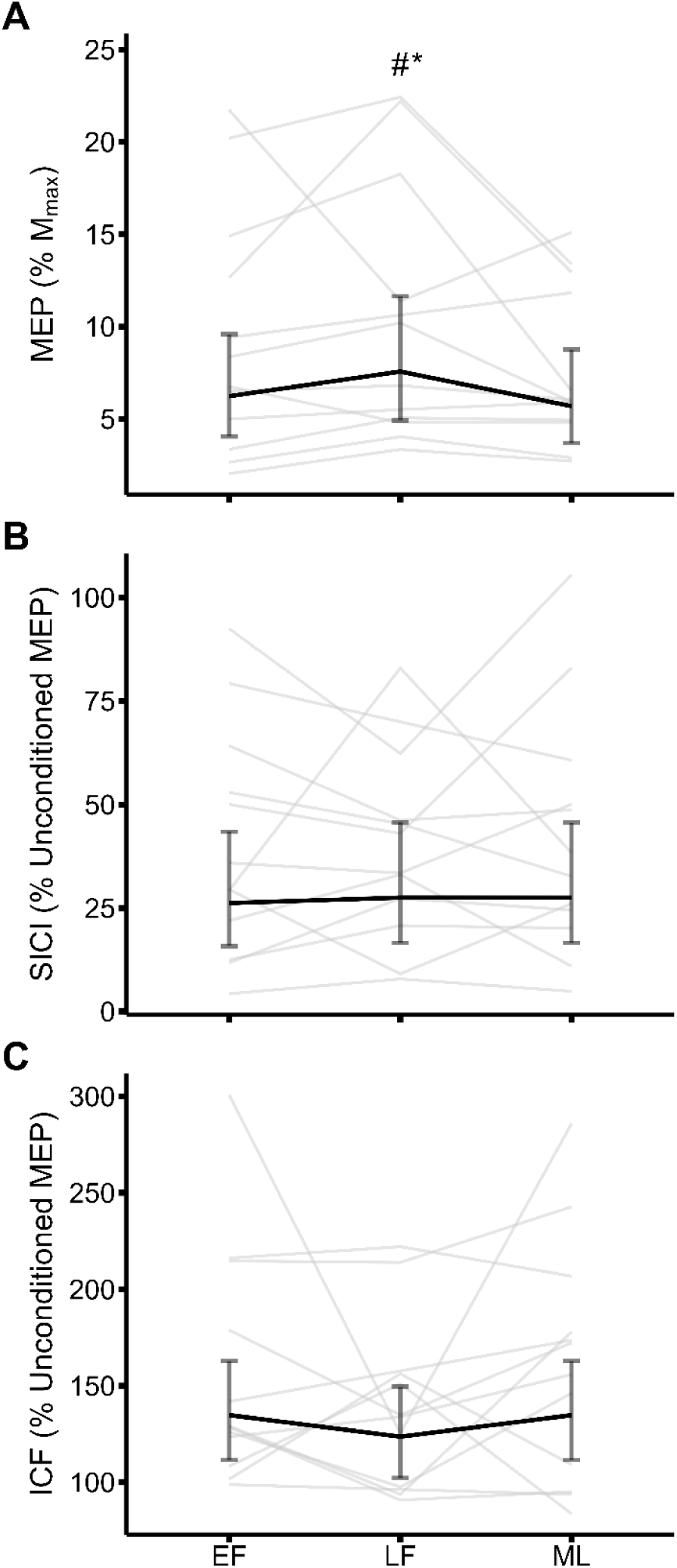
Baseline neurophysiological measures across the menstrual cycle. MEP/M_max_ (A) was modulated by menstrual phase with increased excitability at LF phase. Short-interval cortical inhibition (B) showed no effect of menstrual cycle phase on inhibitory neurotransmission, likewise intracortical facilitation (C) showed no effect of menstrual cycle phase on facilitatory neurotransmission. Bold black lines indicate phase estimated marginal means with error bars indicating 95% confidence intervals. Thin grey lines represent individual participant data. EF: Early Follicular, ICF: Intracortical facilitation, LF: Late Follicular, MEP: Motor evoked potential, ML: Mid-luteal, SICI: Short-interval cortical inhibition, ICF: Intracortical facilitation. * greater than early follicular p <0.05, # greater than mid-luteal p <0.05.

### Spike-timing-dependent plasticity

PAS data, except MEP (% baseline), were logarithmically transformed due to heteroskedastic distribution of the model residuals; back-transformed data are presented in plots for clarity. MEP/M_max_ was significantly affected by phase and time (χ^2^ (4)=12.80, *p* = 0.012), with an interaction between these two factors (χ^2^ (4)=13.46, *p* = 0.009). Whereby, there were no differences between menstrual cycle phases at 15 minutes (*p* ≥ 0.099) but at 30 minutes, ML was greater than both follicular phases (*p* = 0.006 and *p* = 0.048 for EF and LF, respectively). In the EF and LF phases, MEP/M_max_ did not differ from baseline at any time point (*p* ≥ 0.093), although in the LF the MEP/M_max_ decreased from 15 to 30 minutes (7.7 ± 0.6 vs. 6.5 ± 0.5% M_max,_ *p* = 0.015). In the ML phase, MEP/M_max_ did not increase from baseline (6.3 ± 0.4% M_max_) at 15-minutes (6.8 ± 0.5% M_max_, *p* = 0.324), however it increased compared to baseline at 30 minutes (7.5 ± 0.5% M_max_, *p* = 0.004) with no difference between 15 and 30 minutes (*p* = 0.183).

When the MEP response was quantified as a percentage of baseline, it was also influenced by phase and time (χ^2^ (4)=30.57, *p* <0.001) with an interaction between phase and time (χ^2^ (4)=19.94, *p* <0.001). In all phases PAS caused an increase in MEPs at 15 minutes (112 ± 5, 115 ± 5 and 113 ± 7% baseline, *p* ≤ 0.010), but at 30 minutes MEPs returned to baseline in both EF and LF phases (p >0.602), whereas it continued to increase in the ML phase compared to both baseline and 15-minutes (126 ± 7% baseline, *p* <0.001 and *p* = 0.029 respectively).

Following the PAS protocol, SICI showed no effects of phase and time (χ^2^ (4)=0.43, *p* = 0.980). However, ICF was influenced by phase and time (χ^2^ (4)=37.84, *p* <0.001), showing an interaction effect (χ^2^ (4)=15.70, *p* = 0.003). ICF in the EF and ML phases remained constant (*p* ≥0.185), but in the LF phase it increased from baseline (133 ± 7% unconditioned MEP) and 15 minutes (142 ± 8% unconditioned MEP) to 30-minutes (166 ± 9% unconditioned MEP, *p* <0.001 and *p* = 0.006).There was no phase effect at baseline (132 ± 7 and 130 ± 7% unconditioned MEP for EF and ML respectively, *p* >0.948), but the LF phase had greater ICF than EF and ML at 15 (EF 121 ± 6; ML 124 ± 7% unconditioned MEP, *p* <0.014) and 30 minutes (EF 129 ± 7; ML 131 ± 7% unconditioned MEP, *p* <0.001).

## Discussion

In this study we aimed to investigate the effect of fluctuating levels of endogenous female sex hormones across three phases of the menstrual cycle on cortical excitability and neuroplasticity. Contrary to our hypothesis, no baseline changes in inhibitory or facilitatory intracortical neurotransmission were observed, and corticospinal excitability was highest in the LF phase. Furthermore, facilitatory STDP in response to a PAS protocol transiently increased corticospinal excitability regardless of menstrual cycle phase, but the greatest and most sustained magnitude of facilitation was observed in the ML phase, associated with high progesterone and moderate 17β-oestradiol levels.

### Cortical Neuroplasticity

The present findings demonstrate that cortical neuroplasticity can be induced regardless of hormone levels when assessed 15-minutes after a PAS protocol; however, elevated progesterone in the ML phase appears to potentiate and elongate this effect at 30 minutes post-PAS. The present data from the EF phase are in agreement with Tecchio et al. (2008), who tested PAS solely in the EF and reported facilitation of MEPs in the abductor pollicis brevis after 10 minutes. The present data expands upon this by testing over a longer timeframe and across more phases, confirmed with hormone concentrations; however it also contrasts to those using rTMS protocols. Inghilleri et al. (2004) reported a blunting effect on rTMS stimulation trains (5 Hz) on day one of the cycle, compared to MEP facilitation on day fourteen in a high oestrogen state. Ramdeo et al. (2024) measured corticospinal excitability ten minutes after iTBS, with significant facilitation observed in the mid-follicular, which was then blunted in the ML phase. However, as previously mentioned, rTMS protocols stimulate cortical neuroplasticity in a non-selective manner (Player et al., 2012). Indeed, pharmacological studies have found opposing effects in responses to selective vs. non-selective neuroplasticity interventions (Kuo et al., 2008). This suggests that the selective nature of PAS in probing sensorimotor cortical synapses and STDP, may provide specific insights into mechanisms of endogenous hormone level fluctuations across the menstrual cycle on cortical neuroplasticity.

Elevated dopamine (DA) has been shown to prolong the effects of PAS, but this is a non-linear relationship as the prolongation was only evident with moderate dosages (Kuo et al., 2008; Thirugnanasambandam et al., 2011). The potentiation of PAS by DA is largely dependent on NMDA receptor (NMDAr) density (Beaupain et al., 2025), and oestradiol has previously been shown to promote NMDAr density on neuronal dendrites (Woolley et al., 1997). Oestradiol was elevated in the LF and ML phases, presumably increasing NMDAr density in these phases. Progesterone in combination with oestradiol, as seen in the ML phase, has been shown to have no deleterious effect on NMDAr density (Cyr et al., 2000). Progesterone itself has been shown to promote NMDA-induced DA release in the rat striatum (Cabrera & Navarro, 1996). Furthermore, the progesterone metabolite allopregnanolone promotes DA release in some brain regions of the rat (Rouge-Pont et al., 2002). Allopregnanolone increases in tandem with progesterone, although the rate of metabolism in the luteal phase means the magnitude of change is smaller (Kimball et al., 2020). In non-human primates DA receptor availability is greater in the luteal compared to follicular phase (Czoty et al., 2009). Combined this suggests that as oestradiol induces a rise in NMDAr density, that progesterone augments by increasing the release and sensitivity to DA. Therefore, the ML phase likely represents optimal neuro-endocrinological conditions for the peak of the inverted U for DA-mediated PAS prolongation, and also potentially explains the greater and more sustained STDP in the ML phase in the present study (Figure 5).

**Figure 5:**
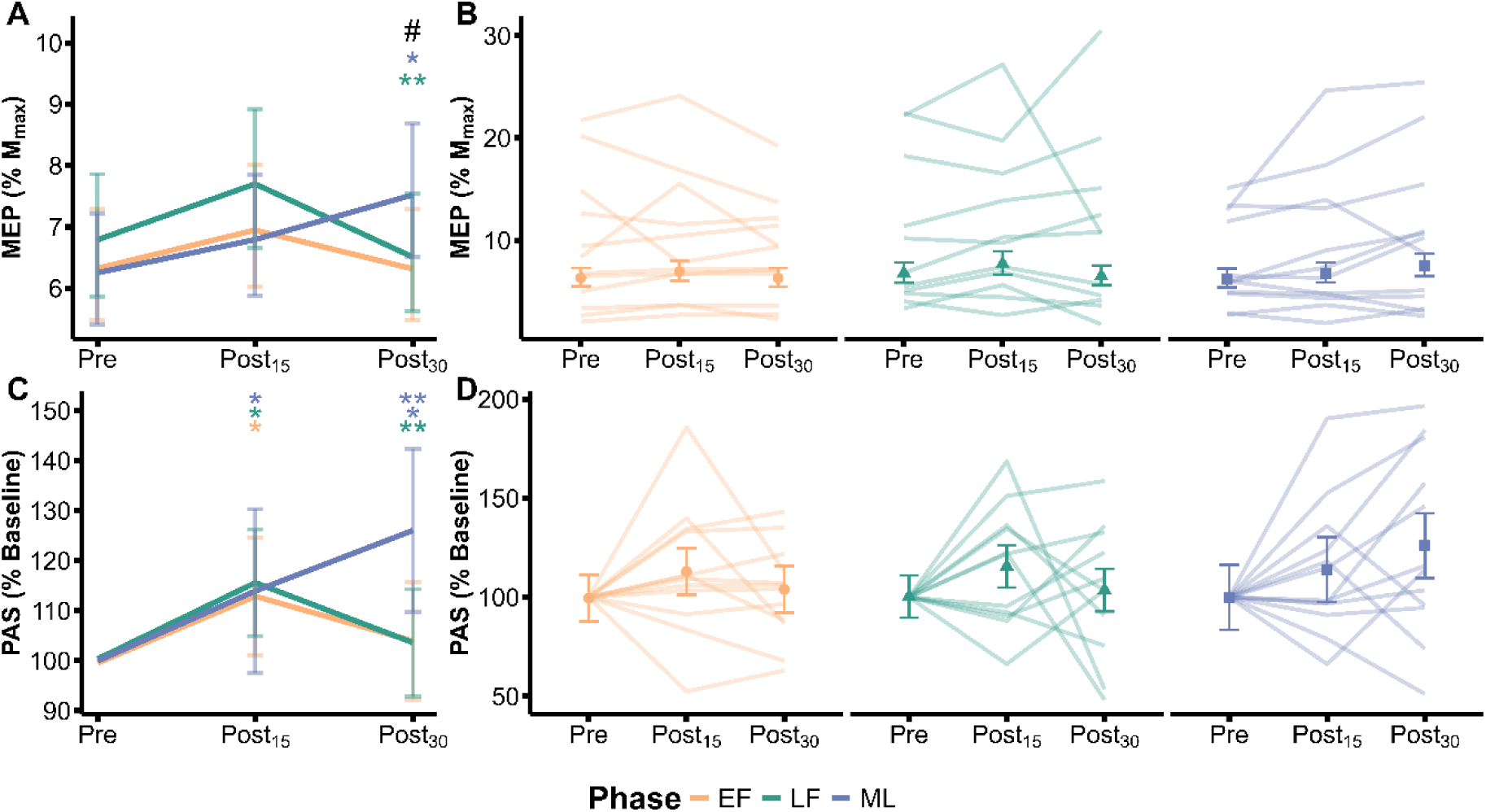
Time course of cortical neuroplasticity following paired associative stimulation. MEP/M_max_ (A) was modulated time and menstrual cycle phase. When expressed as percentage change (C) ML phase showed continued and greater magnitude of neuroplasticity response. The individual responses by phase for both are presented in panel B and D respectively. Bold lines indicate phase estimated marginal means with error bars indicating 95% confidence intervals. Thin lines represent individual participant data. EF: Early Follicular, LF: Late Follicular, MEP: Motor evoked potential, ML: Mid-luteal, PAS: Paired associative stimulation. # mid-luteal phase greater than early and late follicular, * significantly greater than Pre, ** significantly different than Post_15_.

**Figure 6:**
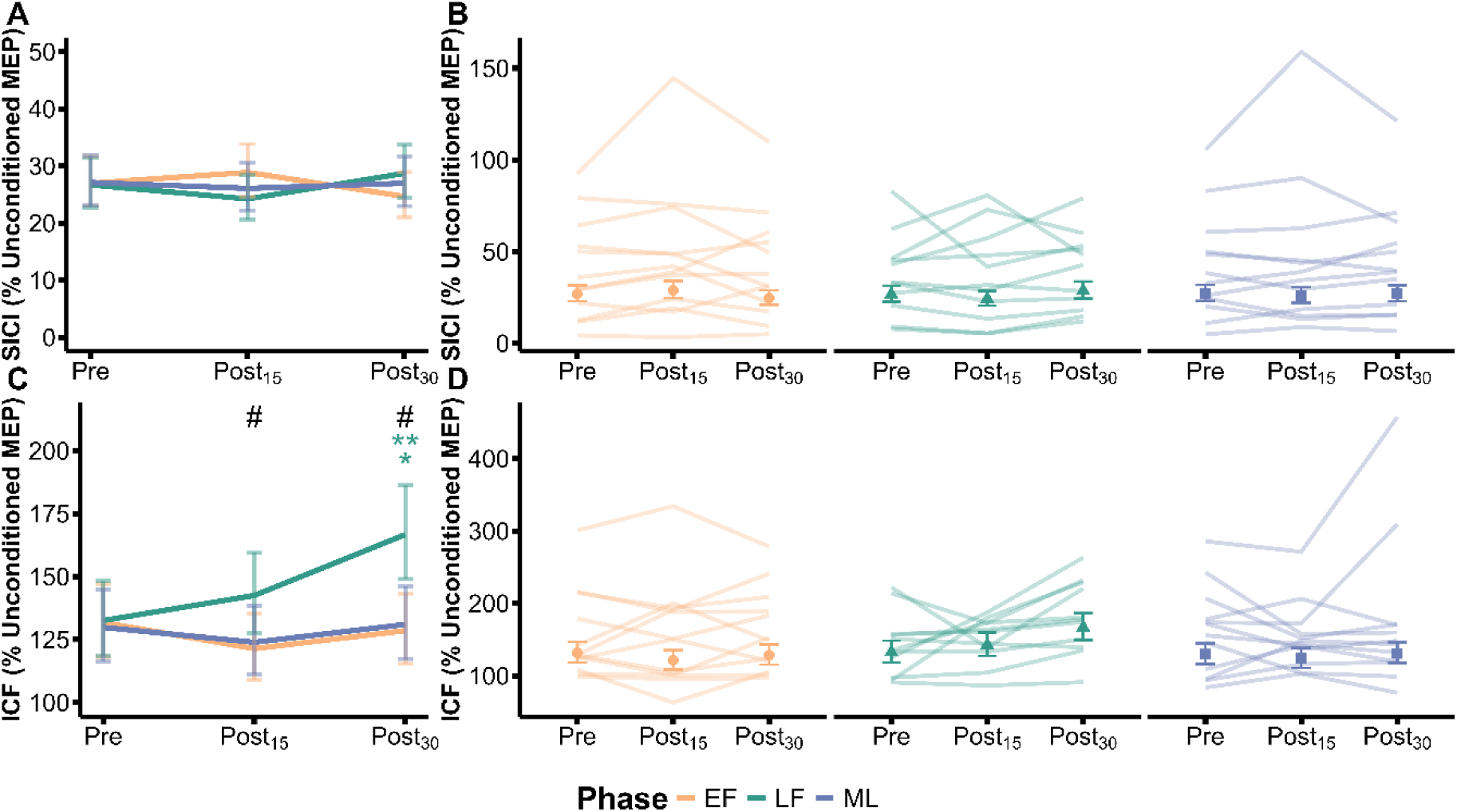
Time course of cortical neurotransmission changes following paired associative stimulation. Short intracortical inhibition (A) showed no changes after PAS. Individual responses for short intracortical inhibition across phase are shown in panel B. However, intracortical facilitation (C) increased in the LF phase only after the PAS protocol. The individual responses (D) illustrate the consistency of this in the late follicular phase. Bold lines indicate phase estimated marginal means with error bars indicating 95% confidence intervals. Thin lines represent individual participant data. EF: Early Follicular, LF: Late Follicular, MEP: Motor evoked potential, ML: Mid-luteal, PAS: Paired associative stimulation. # late follicular phase higher than early follicular and mid-luteal, * significantly greater than Pre, ** significantly different than Post_15_.

Intracortical GABAergic and glutamatergic neurotransmission do not appear to influence the outcomes of PAS across menstrual cycle phases, as SICI and ICF did not change at baseline, nor was the increased ICF in the LF associated with any PAS induced facilitation. These findings are consistent with previous studies showing no effects of a PAS protocol on SICI (Cirillo et al., 2009; Quartarone et al., 2006; Sale et al., 2007). Whilst the ICF results suggest a specific role for oestradiol in glutamatergic facilitation, it appears that this does not influence the magnitude of MEP facilitation, in agreement with previous reports in mixed sex cohorts (Quartarone et al., 2006; Sale et al., 2007). Collectively, these data indicate that GABAergic/glutamatergic neurotransmission are unlikely to be attributable to the menstrual cycle-related changes in PAS, further supporting the mediating role of dopamine.

### Corticospinal Excitability and Paired Pulse TMS

The increased baseline corticospinal excitability observed alongside an increase in oestradiol in the LF phase is consistent with an excitatory effect of oestradiol on excitatory postsynaptic potentials (EPSPs). In isolated cell preparations, oestradiol increased the amplitude, and to a lesser extent the duration of EPSPs; these non-genomic effects act via non-NMDA post-synaptic glutamate receptors rather than by potentiating glutamate release (Wong & Moss, 1992). This is supported by the lack of change in ICF or SICI, indicating intracortical pre-synaptic facilitation was unlikely to be the basis of the oestrogenic effects. Previously, changes in corticospinal excitability have not been associated with hormone levels (Ansdell et al., 2019; Badawy et al., 2013; Hattemer et al., 2007; Inghilleri et al., 2004; Zoghi et al., 2015). However, this discrepancy may reflect methodological differences, where with the exception of Ansdell et al. (2019), most have utilised rMT as their assessment of corticospinal excitability, or normalised test pulse intensity to produce MEPs of 1 mV (Badawy et al., 2013; Hattemer et al., 2007; Zoghi et al., 2015). As opposed to either input-output curves, or setting a test pulse intensity relative to rMT as used in the present study.

The paired pulse TMS data in a resting hand muscle are consistent with much of the existing literature (Badawy et al., 2013; Hattemer et al., 2007; Zoghi et al., 2015), as there were no observed changes in SICI or ICF across the cycle. This suggests that the potent effects of oestrogen and progesterone seen with *in vitro* preparations, amplifying the excitatory glutamatergic or inhibitory GABAergic inputs (Smith et al., 1987a, 1987b; Smith et al., 1989), are too small to be detected in these muscles at rest. One methodological consideration is the distinction between resting and active assessments of intracortical properties. Ansdell et al. (2019) reported an increase in SICI during a 10% contraction of the knee extensors in the ML phase. It has been shown that a light contraction can reduce MEP variability (Darling et al., 2006), which may partially explain the differences here. Additionally, it could be due to differences in inhibitory projections, as proximal muscles demonstrate lower levels of SICI than distal muscles (Abbruzzese et al., 1999). Nevertheless, one research group has demonstrated greater SICI in hand muscles at rest when progesterone is elevated (Smith et al., 1999), and increased ICF when oestradiol is high (Smith et al., 2002). Notably these studies both normalised conditioning TMS pulses to a percentage of active motor threshold, rather than rMT, and pooled a range of ISIs, rather than the 2 ms used in the present study. Based on the present data, it is likely that the 17β-oestradiol-mediated increases in MEP/M_max_ amplitude reflect alterations in EPSPs, or changes at a sub-cortical level, rather than variations in the balance of intracortical inhibition and facilitation at rest.

### Limitations

One limitation of the present study was participant retention, as detailed in Figure 2, only 43% of participants completed the study with appropriate hormonal phase confirmation. Whilst eleven dropouts were due to normal attrition, importantly five participants were excluded based on cycle irregularities identified through the three-step verification method (Schaumberg et al., 2017). This confirms the value of tracking and hormone phase verification, along with strengthening the conclusions drawn from the final sample. Whilst assessing the real-world feasibility of this type of research is beyond the scope of the present study, these findings illustrate the importance of considering additional dropout when planning similar studies.

A 25 ms ISI between the sensory and motor stimulation was used for all participants during the PAS protocol. Whilst this approach lacks the individualisation of N20 latency-based protocols (Jung et al., 2013), a 25 ms ISI has consistently been shown to induce facilitation (Player et al., 2012; Stefan et al., 2002). As facilitation was observed in at least two-thirds of participants in each phase, this supports previous research on the effectiveness of a 25 ms protocol (Player et al., 2012). Additionally, previous work has shown that individualising the latency does not improve PAS outcomes in some clinical populations (Jung et al., 2013). Therefore, it is unlikely that using a fixed ISI unduly impacted the present study’s aim of investigating changes in facilitatory PAS across the menstrual cycle.

### Conclusions

This study provides new insights into the interaction between female sex hormones and spike-timing-dependent plasticity. The PAS data demonstrates that healthy young eumenorrheic females exhibit neuroplastic capacity regardless of menstrual cycle phase. However elevated progesterone and oestradiol in the ML phase prolongs and potentiates these effects, which has potential implications for motor learning and relearning in rehabilitation contexts. The study also demonstrated that corticospinal excitability is increased in a high-oestradiol, low-progesterone state, while intracortical neurotransmission at rest is unaffected by the menstrual cycle. Importantly, by using rigorous hormonal verification we can begin to understand how study protocols or rehabilitation could be better optimised for females, reinforcing the importance of their continued inclusion in neurophysiological research.

## Acknowledgements

The researchers would like to thank Anna Wigbers and Hannah Wilson for their assistance with aspects of data collection, and the participants of the present study for their time and efforts.

## Data availability statement

The data that support the findings of this study are available from the corresponding author upon reasonable request.

## References

Abbruzzese, G., Assini, A., Buccolieri, A., Schieppati, M., & Trompetto, C. (1999). Comparison of intracortical inhibition and facilitation in distal and proximal arm muscles in humans. J Physiol, 514 (Pt 3)(Pt 3), 895–903. 10.1111/j.1469-7793.1999.895ad.x

Adams, M. M., Fink, S. E., Janssen, W. G., Shah, R. A., & Morrison, J. H. (2004). Estrogen modulates synaptic N-methyl-D-aspartate receptor subunit distribution in the aged hippocampus. J Comp Neurol, 474(3), 419–426. 10.1002/cne.20148

Alder, G., Signal, N., Olsen, S., & Taylor, D. (2019). A Systematic Review of Paired Associative Stimulation (PAS) to Modulate Lower Limb Corticomotor Excitability: Implications for Stimulation Parameter Selection and Experimental Design. Front Neurosci, 13, 895. 10.3389/fnins.2019.00895

Ansdell, P., Brownstein, C. G., Skarabot, J., Hicks, K. M., Simoes, D. C. M., Thomas, K., Howatson, G., Hunter, S. K., & Goodall, S. (2019). Menstrual cycle-associated modulations in neuromuscular function and fatigability of the knee extensors in eumenorrheic women. J Appl Physiol (1985), 126(6), 1701–1712. 10.1152/japplphysiol.01041.2018

Badawy, R. A., Vogrin, S. J., Lai, A., & Cook, M. J. (2013). Are patterns of cortical hyperexcitability altered in catamenial epilepsy? Ann Neurol, 74(5), 743–757. 10.1002/ana.23923

Bates, D., McHler, M., Bolker, B., & Walker, S. (2015). Fitting Linear Mixed-Effects Models Using {lme4}.

Beaupain, M. C., Ghanavati, E., Frese, A. M., Melo, L., Kuo, M. F., & Nitsche, M. A. (2025). NMDA receptor involvement in dopaminergic modulation of neuroplasticity induced by paired associative stimulation. Int J Neuropsychopharmacol, 28(6). 10.1093/ijnp/pyaf038

Brownstein, C. G., Ansdell, P., Skarabot, J., Howatson, G., Goodall, S., & Thomas, K. (2018). An optimal protocol for measurement of corticospinal excitability, short intracortical inhibition and intracortical facilitation in the rectus femoris. J Neurol Sci, 394, 45–56. 10.1016/j.jns.2018.09.001

Cabrera, R. J., & Navarro, C. E. (1996). Progesterone in vitro increases NMDA-evoked [3H] dopamine release from striatal slices in proestrus rats. Neuropharmacology, 35(2), 175–178. 10.1016/0028-3908(95)00152-2

Carson, R. G., & Kennedy, N. C. (2013). Modulation of human corticospinal excitability by paired associative stimulation. Front Hum Neurosci, 7, 823. 10.3389/fnhum.2013.00823

Chinta, P., Rebekah, G., T Kunjummen, A., & S. Kamath, M. (2020). Revisiting the role of serum progesterone as a test of ovulation in eumenorrheic subfertile women: a prospective diagnostic accuracy study. Fertility and Sterility, 114(6), 1315–1321. 10.1016/j.fertnstert.2020.06.030

Cirillo, J., Lavender, A. P., Ridding, M. C., & Semmler, J. G. (2009). Motor cortex plasticity induced by paired associative stimulation is enhanced in physically active individuals. J Physiol, 587(Pt 24), 5831–5842. 10.1113/jphysiol.2009.181834

Cyr, M., Ghribi, O., & Di Paolo, T. (2000). Regional and selective effects of oestradiol and progesterone on NMDA and AMPA receptors in the rat brain. J Neuroendocrinol, 12(5), 445–452. 10.1046/j.1365-2826.2000.00471.x

Czoty, P. W., Riddick, N. V., Gage, H. D., Sandridge, M., Nader, S. H., Garg, S., Bounds, M., Garg, P. K., & Nader, M. A. (2009). Effect of menstrual cycle phase on dopamine D2 receptor availability in female cynomolgus monkeys. Neuropsychopharmacology, 34(3), 548–554. 10.1038/npp.2008.3

Darling, W. G., Wolf, S. L., & Butler, A. J. (2006). Variability of motor potentials evoked by transcranial magnetic stimulation depends on muscle activation. Exp Brain Res, 174(2), 376–385. 10.1007/s00221-006-0468-9

Di Lazzaro, V., Pilato, F., Dileone, M., Profice, P., Ranieri, F., Ricci, V., Bria, P., Tonali, P. A., & Ziemann, U. (2007). Segregating two inhibitory circuits in human motor cortex at the level of GABAA receptor subtypes: a TMS study. Clin Neurophysiol, 118(10), 2207–2214. 10.1016/j.clinph.2007.07.005

El-Sayes, J., Turco, C. V., Skelly, L. E., Nicolini, C., Fahnestock, M., Gibala, M. J., & Nelson, A. J. (2019). The Effects of Biological Sex and Ovarian Hormones on Exercise-Induced Neuroplasticity. Neuroscience, 410, 29–40. 10.1016/j.neuroscience.2019.04.054

Grover, F. M., Chen, B., & Perez, M. A. (2023). Increased paired stimuli enhance corticospinal-motoneuronal plasticity in humans with spinal cord injury. J Neurophysiol, 129(6), 1414–1422. 10.1152/jn.00499.2022

Hattemer, K., Knake, S., Reis, J., Rochon, J., Oertel, W. H., Rosenow, F., & Hamer, H. M. (2007). Excitability of the motor cortex during ovulatory and anovulatory cycles: a transcranial magnetic stimulation study. Clin Endocrinol (Oxf), 66(3), 387–393. 10.1111/j.1365-2265.2007.02744.x

Inghilleri, M., Conte, A., Curra, A., Frasca, V., Lorenzano, C., & Berardelli, A. (2004). Ovarian hormones and cortical excitability. An rTMS study in humans. Clin Neurophysiol, 115(5), 1063–1068. 10.1016/j.clinph.2003.12.003

Jannati, A., Oberman, L. M., Rotenberg, A., & Pascual-Leone, A. (2022). Assessing the mechanisms of brain plasticity by transcranial magnetic stimulation. Neuropsychopharmacology. 10.1038/s41386-022-01453-8

Jenz, S. T., & Pearcey, G. E. P. (2022). Sex matters in neuromuscular control. Acta Physiol (Oxf), 235(2), e13823. 10.1111/apha.13823

Jung, N. H., Janzarik, W. G., Delvendahl, I., Munchau, A., Biscaldi, M., Mainberger, F., Baumer, T., Rauh, R., & Mall, V. (2013). Impaired induction of long-term potentiation-like plasticity in patients with high-functioning autism and Asperger syndrome. Dev Med Child Neurol, 55(1), 83–89. 10.1111/dmcn.12012

Jung, P., & Ziemann, U. (2009). Homeostatic and nonhomeostatic modulation of learning in human motor cortex. Journal of Neuroscience, 29(17), 5597–5604. 10.1523/JNEUROSCI.0222-09.2009

Kimball, A., Dichtel, L. E., Nyer, M. B., Mischoulon, D., Fisher, L. B., Cusin, C., Dording, C. M., Trinh, N. H., Yeung, A., Haines, M. S., Sung, J. C., Pinna, G., Rasmusson, A. M., Carpenter, L. L., Fava, M., Klibanski, A., & Miller, K. K. (2020). The allopregnanolone to progesterone ratio across the menstrual cycle and in menopause. Psychoneuroendocrinology, 112, 104512. 10.1016/j.psyneuen.2019.104512

Kujirai, T., Caramia, M. D., Rothwell, J. C., Day, B. L., Thompson, P. D., Ferbert, A., Wroe, S., Asselman, P., & Marsden, C. D. (1993). Corticocortical inhibition in human motor cortex. J Physiol, 471, 501–519. 10.1113/jphysiol.1993.sp019912

Kuo, M. F., Paulus, W., & Nitsche, M. A. (2008). Boosting focally-induced brain plasticity by dopamine. Cereb Cortex, 18(3), 648–651. 10.1093/cercor/bhm098

Kuznetsova, A., Brockhoff, P., Brockhoff, B., Christensen, R., Christensen, H., & Christensen, B. (2017). {lmerTest} Package: Tests in Linear Mixed Effects Models.

Lenth, R. V. (2025). emmeans: Estimated Marginal Means, aka Least-Squares Means. In. Maeda, F., Keenan, J. P., Tormos, J. M., Topka, H., & Pascual-Leone, A. (2000). Interindividual variability of the modulatory effects of repetitive transcranial magnetic stimulation on cortical excitability. Exp Brain Res, 133(4), 425–430. 10.1007/s002210000432

Malcolm, C. E., & Cumming, D. C. (2003). Does anovulation exist in eumenorrheic women? Obstet Gynecol, 102(2), 317–318. 10.1016/s0029-7844(03)00527-1

Oldfield, R. C. (1971). The assessment and analysis of handedness: the Edinburgh inventory. Neuropsychologia, 9(1), 97–113. 10.1016/0028-3932(71)90067-4

ONS, Office for National Statistics (2023). Population estimates for the UK, England, Wales, Scotland, and Northern Ireland: mid-2023.

Pieber, K., Herceg, M., Paternostro-Sluga, T., & Schuhfried, O. (2015). Optimizing stimulation parameters in functional electrical stimulation of denervated muscles: a cross-sectional study. Journal of NeuroEngineering and Rehabilitation, 12(1), 51. 10.1186/s12984-015-0046-0

Player, M. J., Taylor, J. L., Alonzo, A., & Loo, C. K. (2012). Paired associative stimulation increases motor cortex excitability more effectively than theta-burst stimulation. Clin Neurophysiol, 123(11), 2220–2226. 10.1016/j.clinph.2012.03.081

Polimanti, R., Simonelli, I., Zappasodi, F., Ventriglia, M., Pellicciari, M. C., Benussi, L., Squitti, R., Rossini, P. M., & Tecchio, F. (2016). Biological factors and age-dependence of primary motor cortex experimental plasticity. Neurol Sci, 37(2), 211–218. 10.1007/s10072-015-2388-6

Quartarone, A., Rizzo, V., Bagnato, S., Morgante, F., Sant’Angelo, A., Girlanda, P., & Siebner, H. R. (2006). Rapid-rate paired associative stimulation of the median nerve and motor cortex can produce long-lasting changes in motor cortical excitability in humans. J Physiol, 575(Pt 2), 657–670. 10.1113/jphysiol.2006.114025

R Core Team. (2024). R: A Language and Environment for Statistical Computing. In (Version 4.4.1) R Foundation for Statistical Computing. https://www.R-project.org

Ramdeo, K. R., Adams, F. C., Drapeau, C. C., Foglia, S. D., Cuizon, M. C., Sader, M. A., Nucci, R., & Nelson, A. J. (2024). The influence of menstrual phase on synaptic plasticity induced via intermittent theta-burst stimulation. Neuroscience, 558, 122–127. 10.1016/j.neuroscience.2024.08.023

Rossi, S., Hallett, M., Rossini, P. M., & Pascual-Leone, A. (2011). Screening questionnaire before TMS: an update. Clin Neurophysiol, 122(8), 1686. 10.1016/j.clinph.2010.12.037

Rossini, P. M., Burke, D., Chen, R., Cohen, L. G., Daskalakis, Z., Di Iorio, R., Di Lazzaro, V., Ferreri, F., Fitzgerald, P. B., George, M. S., Hallett, M., Lefaucheur, J. P., Langguth, B., Matsumoto, H., Miniussi, C., Nitsche, M. A., Pascual-Leone, A., Paulus, W., Rossi, S., . . . Ziemann, U. (2015). Non-invasive electrical and magnetic stimulation of the brain, spinal cord, roots and peripheral nerves: Basic principles and procedures for routine clinical and research application. An updated report from an I.F.C.N. Committee. Clin Neurophysiol, 126(6), 1071–1107. 10.1016/j.clinph.2015.02.001

Rouge-Pont, F., Mayo, W., Marinelli, M., Gingras, M., Le Moal, M., & Piazza, P. V. (2002). The neurosteroid allopregnanolone increases dopamine release and dopaminergic response to morphine in the rat nucleus accumbens. Eur J Neurosci, 16(1), 169–173. 10.1046/j.1460-9568.2002.02084.x

Sale, M. V., Ridding, M. C., & Nordstrom, M. A. (2007). Factors influencing the magnitude and reproducibility of corticomotor excitability changes induced by paired associative stimulation. Exp Brain Res, 181(4), 615–626. 10.1007/s00221-007-0960-x

Schaumberg, M. A., Jenkins, D. G., Janse de Jonge, X. A. K., Emmerton, L. M., & Skinner, T. L. (2017). Three-step method for menstrual and oral contraceptive cycle verification. J Sci Med Sport, 20(11), 965–969. 10.1016/j.jsams.2016.08.013

Smith, C. C., & McMahon, L. L. (2005). Estrogen-induced increase in the magnitude of long-term potentiation occurs only when the ratio of NMDA transmission to AMPA transmission is increased. Journal of Neuroscience, 25(34), 7780–7791. 10.1523/JNEUROSCI.0762-05.2005

Smith, M. J., Adams, L. F., Schmidt, P. J., Rubinow, D. R., & Wassermann, E. M. (2002). Effects of ovarian hormones on human cortical excitability. Ann Neurol, 51(5), 599–603. 10.1002/ana.10180

Smith, M. J., Adams, L. F., Schmidt, P. J., Rubinow, D. R., & Wassermann, E. M. (2003). Abnormal luteal phase excitability of the motor cortex in women with premenstrual syndrome. Biol Psychiatry, 54(7), 757–762. 10.1016/s0006-3223(02)01924-8

Smith, M. J., Keel, J. C., Greenberg, B. D., Adams, L. F., Schmidt, P. J., Rubinow, D. A., & Wassermann, E. M. (1999). Menstrual cycle effects on cortical excitability. Neurology, 53(9), 2069–2069. 10.1212/wnl.53.9.2069

Smith, S. S., Waterhouse, B. D., & Woodward, D. J. (1987a). Locally applied progesterone metabolites alter neuronal responsiveness in the cerebellum. Brain Res Bull, 18(6), 739–747. 10.1016/0361-9230(87)90209-7

Smith, S. S., Waterhouse, B. D., & Woodward, D. J. (1987b). Sex steroid effects on extrahypothalamic CNS. I. Estrogen augments neuronal responsiveness to iontophoretically applied glutamate in the cerebellum. Brain Res, 422(1), 40–51. 10.1016/0006-8993(87)90538-5

Smith, S. S., Woodward, D. J., & Chapin, J. K. (1989). Sex steroids modulate motor-correlated increases in cerebellar discharge. Brain Res, 476(2), 307–316. 10.1016/0006-8993(89)91251-1

Stefan, K., Kunesch, E., Benecke, R., Cohen, L. G., & Classen, J. (2002). Mechanisms of enhancement of human motor cortex excitability induced by interventional paired associative stimulation. J Physiol, 543(Pt 2), 699–708. 10.1113/jphysiol.2002.023317

Stefan, K., Kunesch, E., Cohen, L. G., Benecke, R., & Classen, J. (2000). Induction of plasticity in the human motor cortex by paired associative stimulation [Article]. Brain, 123(3), 572–584. 10.1093/brain/123.3.572

Suppa, A., Quartarone, A., Siebner, H., Chen, R., Di Lazzaro, V., Del Giudice, P., Paulus, W., Rothwell, J. C., Ziemann, U., & Classen, J. (2017). The associative brain at work: Evidence from paired associative stimulation studies in humans. Clin Neurophysiol, 128(11), 2140–2164. 10.1016/j.clinph.2017.08.003

Tecchio, F., Zappasodi, F., Pasqualetti, P., Gennaro, L., Pellicciari, M. C., Ercolani, M., Squitti, R., & Rossini, P. M. (2008). Age dependence of primary motor cortex plasticity induced by paired associative stimulation. Clin Neurophysiol, 119(3), 675–682. 10.1016/j.clinph.2007.10.023

Thirugnanasambandam, N., Grundey, J., Paulus, W., & Nitsche, M. A. (2011). Dose-dependent nonlinear effect of L-DOPA on paired associative stimulation-induced neuroplasticity in humans. Journal of Neuroscience, 31(14), 5294–5299. 10.1523/JNEUROSCI.6258-10.2011

Thomas, K., Elmeua, M., Howatson, G., & Goodall, S. (2016). Intensity-Dependent Contribution of Neuromuscular Fatigue after Constant-Load Cycling. Medicine & Science in Sports & Exercise, 48(9), 1751–1760. 10.1249/mss.0000000000000950

Wathen, N. C., Perry, L., Lilford, R. J., & Chard, T. (1984). Interpretation of single progesterone measurement in diagnosis of anovulation and defective luteal phase: observations on analysis of the normal range. Br Med J (Clin Res Ed), 288(6410), 7–9. 10.1136/bmj.288.6410.7

Wickham, H. (2016). ggplot2: Elegant Graphics for Data Analysis. Springer-Verlag New York. https://ggplot2.tidyverse.org

Wischnewski, M., & Schutter, D. (2016). Efficacy and time course of paired associative stimulation in cortical plasticity: Implications for neuropsychiatry. Clin Neurophysiol, 127(1), 732–739. 10.1016/j.clinph.2015.04.072

Woitowich, N. C., Beery, A., & Woodruff, T. (2020). A 10-year follow-up study of sex inclusion in the biological sciences. Elife, 9. 10.7554/eLife.56344

Wong, M., & Moss, R. L. (1992). Long-term and short-term electrophysiological effects of estrogen on the synaptic properties of hippocampal CA1 neurons. Journal of Neuroscience, 12(8), 3217–3225. 10.1523/JNEUROSCI.12-08-03217.1992

Woolley, C. S., Weiland, N. G., McEwen, B. S., & Schwartzkroin, P. A. (1997). Estradiol increases the sensitivity of hippocampal CA1 pyramidal cells to NMDA receptor-mediated synaptic input: correlation with dendritic spine density. Journal of Neuroscience, 17(5), 1848–1859. 10.1523/JNEUROSCI.17-05-01848.1997

Zoghi, M., Vaseghi, B., Bastani, A., Jaberzadeh, S., & Galea, M. P. (2015). The Effects of Sex Hormonal Fluctuations during Menstrual Cycle on Cortical Excitability and Manual Dexterity (a Pilot Study). PLoS One, 10(8), e0136081. 10.1371/journal.pone.0136081

